# The Impact of Malaria-Induced Neutrophil Subset Shift and a Link to Burkitt Lymphoma

**DOI:** 10.1101/2025.10.13.681992

**Authors:** Sharon Akinyi, Ronald K. Tonui, Titus K. Maina, Eddy Agwati, Cliff I. Oduor, Festus M. Njuguna, Kibet K. Keitany, Daniel Chepsiror, Cyrus Ayieko, Ann Moormann, Ann W. Kinyua, Catherine S. Forconi

## Abstract

Burkitt lymphoma (BL) is an aggressive B-cell lymphoma that remains a leading cause of childhood cancer mortality in sub-Saharan Africa. Although the epidemiological link between *Plasmodium falciparum (Pf)* malaria and BL has been established, our understanding of the underlying immunological mechanisms conducive to tumorigenesis is incomplete. To address a noted gap in our knowledge of the immune landscape, we profiled neutrophil subsets from children with different exposure histories to *Pf*-malaria and children diagnosed with BL from Western Kenya, along with healthy malaria-naive Kenyan adults. Using multiparameter flow cytometry, we characterized neutrophils by expression of CD15, CD16, CD10, CD11b, CD182, CD184, and CD62L and found that malaria-exposed children exhibited increased frequencies of aged neutrophil subsets, accompanied by a reduction in the mature subset frequencies compared to malaria-naive children. Malaria-exposed children also had neutrophil profiles that closely resembled those seen in the adults. Notably, a positive correlation (rs *= 0*.*7; p < 0*.*0001)* was observed in immature neutrophils between malaria-exposed healthy and BL children, indicating a similar expansion pattern of this subset in both groups. This finding suggests a malaria-driven expansion of the immature subset, potentially promoting a permissive environment for BL. Our data suggests that the observed shift in neutrophil profiles could contribute to the malaria-induced immunopathology associated with BL

**Visual abstract:** Created in BioRender. Forconi, C. (2025) https://BioRender.com/oz60qvq

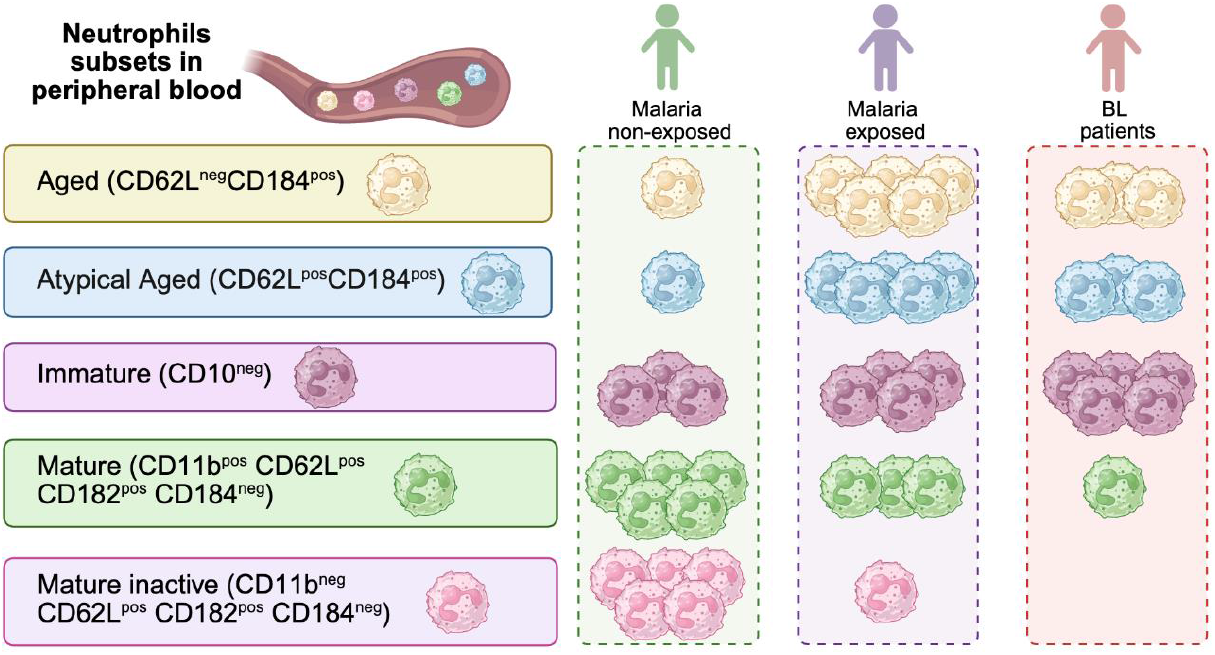

## Introduction

Endemic Burkitt lymphoma (BL) is an aggressive childhood B-cell lymphoma that accounts for 50%–70% of childhood cancers in sub-Saharan Africa (1,2), where malaria is also endemic (3). Chronic malaria exposure predisposes children to BL by inducing defects in host immune responses (4). The defects present as reduced IFN-γ T cell responses, suppression of T cell responses, as well as a skewed T cell immune profile (5,6,7). The result of these alterations is the impairment of immune surveillance against Epstein-Barr Virus (EBV), a well-established oncogenic cofactor in eBL pathogenesis, leading to loss of viral control and thereby contributing to BL oncogenesis (8). While most research has focused on adaptive immunity, malaria-induced modifications in innate immune cells have been explored less extensively, yet these modifications may also play an important role in BL pathogenesis.

Neutrophils, the most abundant innate immune cell in human circulation (9,10), are rapidly recruited to sites of infection, where they deploy diverse effector functions, including the release of neutrophil extracellular traps (NETs), to limit viral replication and eliminate pathogens (11,12). Beyond direct pathogen clearance, they have antigen-presenting capacity, allowing them to activate, shape, and regulate antiviral adaptive immune responses (13). Neutrophils are increasingly being recognized as a heterogeneous population whose subsets dynamically adapt to physiological and pathological conditions (14). However, extensive exploration of neutrophil biology in the context of chronic malaria has been limited. This is largely due to their short lifespan, limited survival after cryopreservation, and logistical challenges of performing real-time flow cytometric analyses in resource-limited settings typical of sub-Saharan Africa. As a result, the contribution of malaria-driven neutrophil reprogramming to BL pathogenesis remains largely unexplored.

Inflammatory cues drive neutrophil plasticity (15,16,17), promoting the release of immature subset from bone marrow and retention of aged subset in circulation (18,19). Persistent exposure to inflammation induces metabolic and transcriptional reprogramming, leading to adaptations that last for weeks to several months (20). *Plasmodium falciparum (Pf)* malaria is a highly inflammatory disease (21) that is known to impair neutrophil effector functions (22,23,24), as well as impair neutrophil activation in children (25). In malaria endemic regions, where chronic *Pf* exposure is common, chronic immune stimulation may induce long-term transcriptional reprogramming of the neutrophil neutrotime, therefore altering circulating neutrophil subsets in ways that are not fully understood. It therefore remains unknown how chronic *Pf* malaria infection influences neutrophil subset dynamics and whether these alterations influence BL development and pathogenesis.

This work aimed to ascertain the effects of chronic *Pf*-malaria exposure on neutrophil subset composition and evaluate their association with BL. Our study characterized neutrophil subsets from the peripheral blood of malaria-exposed, malaria-naive, and BL children. Aged neutrophils are typically defined by the expression of the aging marker CD184 and loss of the adhesion molecule CD62L, which facilitates their migration to the site of inflammation. In addition to the conventional aged neutrophil subset (CD62L^neg^CD184^pos^), we identified an additional subset expressing both the aging marker CD184 and the adhesion molecule CD62L, and labelled it “atypical aged” (CD62L^pos^CD184^pos^). This subset was consistent in all our participants and, to our knowledge, has not been previously reported, suggesting increased heterogeneity in the aged neutrophil subset.

## Materials and Methods

### Participants

To interpret malaria-induced alterations in neutrophil subsets, we enrolled malaria-naive healthy children to establish a baseline distribution and healthy adults to represent neutrophil profiles of a fully mature immune system. Children with non-BL cancers were also included to provide insight into neutrophil profiles in other pediatric malignancies. We enrolled participants within the age range of BL development, 4–9 years (26), 19 healthy children residing in a malaria holoendemic area (malaria-exposed), 11 healthy children residing in the low malaria transmission area (malaria-naive), 11 children diagnosed with BL, and 5 children diagnosed with non-BL cancers, such as Hodgkin’s lymphomas, non-Hodgkin lymphomas, and sarcomas. We also included 3 Kenyan adult healthy controls. All the cancer cases were enrolled at the Moi Teaching and Referral Hospital, Eldoret, Kenya. The cancer diagnosis was made by flow cytometry, and the hospital pathologist confirmed the diagnosis by histology (hematoxylin and eosin tissue staining). The malaria-exposed healthy controls were enrolled at Ahero subcounty hospital, a region known for high *Pf* malaria transmission as well as high morbidity and mortality cases resulting from *Pf* malaria (27) in Kisumu County, Kenya. The malaria-naive healthy controls and healthy adults were enrolled in Mosoriot hospital, Nandi County, Kenya, a region with seasonal malaria transmission with a considerable year-to-year variation (28) known as a malaria hypoendemic area.

### Neutrophil Immunophenotyping

For the identification and characterization of neutrophils, 100 μL of whole blood was collected in lithium heparin tubes, and a nine-color panel was used to identify and characterize neutrophil subsets by flow cytometry. The antibodies in the panel are summarised in S1 TABLE. After staining, samples were fixed with 1X fixation and Permeabilization Solution containing 4.2% formaldehyde (BD Biosciences #Cat:51-2090KZ). After fixing, the samples were stored at 4°C and analyzed using the Beckman Coulter CytoFLEX Flow Cytometer the following day. Compensation controls and FMOs were prepared for each run (*Full protocol detail in Supporting information)*. Using FlowJo software, version 10.10.0, the compensation controls were used to correct for fluorescence spillovers, and the FMOs were used to set gates. A flow cytometry gating strategy was used for phenotyping neutrophils, and Boolean gating was used to identify neutrophil subsets **S1 FIGURE A**. The CD62L FMO and gating strategy used to identify the aged and atypical aged subsets are shown in **S1 FIGURE B** and **S1 FIGURE C**, respectively.

### Serological Testing for AMA1 Antibody Levels

To confirm malaria exposure history, antibody levels against Apical Membrane Antigen 1 (AMA1, gift from Dr. Dutta, Walter Reed Army Institute of Research) were measured along with beads coupled to bovine serum albumine (BSA, Millipore Sigma) as internal control. Briefly, 50μl of AMA1 and BSA conjugated beads (500 beads of each per well), suspended in assay buffer (PBS pH7.4 containing 0.1% BSA, 0.05% Tween-20, and 0.05% sodium azide) was aliquoted into the appropriately labelled wells of a flat bottom uncoated 96-well plate and a magnetic bead washer was then used to wash the beads in plain ABE for 2 minutes. 50μl of 1/100 diluted human plasma was added to the respective wells containing beads. A pool of plasma from healthy African adults was used as a standard (2-fold dilution, S1 to S7) while the plasma from confirmed malaria-naive American adults was used as a negative control (dilution 1/100). The plate was incubated at room temperature for 2 hours. The beads were washed 3 times in the assay buffer and incubated for another 1 hour at room temperature in 50μl of biotinylated anti-human IgG. The washing of the beads was repeated three times, followed by a 15-minute incubation at room temperature with phycoerythrin-conjugated streptavidin. After 15 minutes, the plate was washed for the final 3 times, and was read on a Flex Map 3D analyzer. A minimum of 50 beads per analyte was acquired per well. The level of specific anti-AMA1 IgG from each of our studied groups was determined after subtraction of the BSA-beads median fluorescence intensity (MFI) to take into account possible non-specific binding.

### DNA extraction and malaria detection by qPCR

Genomic DNA was extracted from 100 μL of whole blood using the Zymo Quick DNA 96 kit (Zymo Research D3012) following the manufacturer’s instructions. Briefly, blood samples were lysed in 400 μL Genomic Lysis Buffer before the mixture was transferred to a Silicon-A Plate on a Collection Plate and centrifuged at 3800 g (Ependorf Centrifuge 5810) for 5 minutes. To obtain pure DNA, the extract was cleaned in 200 uL DNA Pre-Wash buffer followed by g-DNA Wash buffer, spnning each wash at 3800gs for 5 minutes and discarding the supernatants. The DNA was eluted in 30uL of the DNA Elution Buffer and stored at -20^°^C for downstream analysis. The quality of the DNA product was confirmed using spectrophotometry (Thermo Scientific Nanodrop 2000) at A260/A280 and A260/A230.

For malaria parasite detection, we utlized an ultrasensitive quantitative real time polymeraase chain reaction (qPCR) based on the varATS region using foward primer, reverse primer and probe sequences originally described in (29). To prepare 12.5uL reaction mix, 4.75 uL of the DNA template was added to 7.75 uL of the qPCR mastermix prepared from 6.25 uL TaqMan Gene Expression Mastermix (Thermo Fisher 43690116), 0.5uL 20 uM Fw (IDT), 0.5uL 20 uM Rev (IDT), 0.25 uL 20 uM (IDT) and 0.5 uL Molecular grade Water (Thermo Fisher 10977035). The 12.5 μL reaction mix was loaded into a 0.1 mL MicroAmp Fast 96-Well Reaction plate (Thermo Fisher 4346907), sealed with MicroAmp Optical Adhesive Film (Thermo Fisher 431197), and loaded on QuantStudio 6 pro qPCR machine (Thermo Fisher A43180). The qPCR run protocol was set for 2 minutes pre-incubation at 50 °C, 10 minutes initial denaturation at 95 °C, and 45 cycles of 15 seconds denaturation at 95 °C followed by 1 minute annealing and elongation steps at 55 °C. The qPCR output was analysed using Design and Analysis Software version 2.6 QuantStudio 6/7 Pro Systems (Thermo Fisher). For quality control, the samples were run alongside replicates of positive and negative controls. To determine parasite densities, we conducted a standard curve analysis using a 10-fold serial dilution of Pf positive control (10,000, 1,000, 100, 10, 1, and 0.1 copies per μL).

### Statistical analysis

The flow cytometry data was analyzed using FlowJo version 10.10.0. Statistical analysis was done in R software version 4.2.1. The data was not normally distributed, so the Mann-Whitneytest was used to compare subset frequency between the two groups. For comparisons involving more than two groups, the Kruskal-Wallis test was applied to detect overall differences, followed by Dunn’s post hoc test, allowing for Dunn’s multiple comparison correction. Since the data was not uniformly distributed, the Spearman correlation was used. A predetermined *p*-value < 0.05 was considered statistically significant.

### Ethics statement

Ethical approval was obtained from the Scientific and Ethical Review Unit (SERU) at the Kenya Medical Research Institute (KEMRI), the University of Massachusetts Chan (UMass Chan) Medical School, and the Institutional Research Ethics Committee (IREC) at Moi Teaching and Referral Hospital/Moi University College of Health Sciences. Parents of participants under the age of 10 years provided written informed consent, while participants aged 10-17 years provided signed assent in addition to the parent’s written informed consent.

## Results

### Characteristics of the study participants

As expected, children with cancer had significantly higher Absolute Neutrophil Count (ANC, *p=0*.*0001, p=0*.*02*) and White Blood Cell Count (WBC, *p=0*.*01, p=0*.*03*) compared to healthy control children. Children with other childhood cancers showed significantly higher ANC counts compared to BL children (*p=*0.04), data in **S2 TABLE**. ANC and WBC were comparable between malaria-exposed and malaria-naive healthy children, data in **S2 TABLE**, respectively. We found that 68% of the malaria-exposed healthy controls had asymptomatic malaria infections. As expected, malaria-exposed children had significantly higher AMA1 antibody levels than malaria-naive children: (median MFI 72126, range [384-217702] vs median MFI 0, range [0-2541]; *p=0*.*0001*). There was no significant difference in AMA1 antibody levels between malaria-exposed children and children with BL (median MFI 72126, range [384-217702] vs median MFI 2495, range [0-171811]; *p=0*.*10*), as well as between children with BL and children with other cancers (median MFI 2495, range [0-171811] vs median MFI 197, range [0-153896]; *p=0*.*27*) data in **S2 TABLE**. The demographic data for healthy Kenyan adults is summarized in **S3 TABLE**.

### Malaria shifts “mature” neutrophils subsets towards “aged” phenotype

We characterized and compared the frequency of neutrophil subsets in malaria-exposed (ME) and malaria-naive (MN) children and found that MN children had significantly higher frequencies of mature neutrophils compared to ME (MN median 31%, range [16.4%–56.3%] vs. ME median 2.3%, range [0%–4.97%]; *p=0*.*0017*, respectively **Fig. 1A**). Similar observation was made regarding the frequencies of mature inactive neutrophil (median 0% vs. median 0.08%, range [0–0.37]; *p<0*.*0001*, respectively **Fig. 1B**). No statistical difference was found in regards to immature neutrophils between MN and ME (**Fig. 1C**), however, we noticed significantly high frequencies of atypical aged and aged neutrophils in ME children compared to MN ones with a median of 35.8% atypical aged neutrophils in ME, range [11.7%–49.3%] vs. median 0.02% in MN, range [0.02%–0.64%] *p<0*.*0001*, respectively, **Fig. 1D;** and a median of 9.35% aged neutrophils in ME, (range [3.2%–34.6%]) vs. a median of 0.3% in MN (range [0.15%–0.78%]), *p<0*.*0001*, **Fig.1E**). Together this suggests that malaria exposure is driving the expansion of the aged neutrophils.

**Figure 1:**
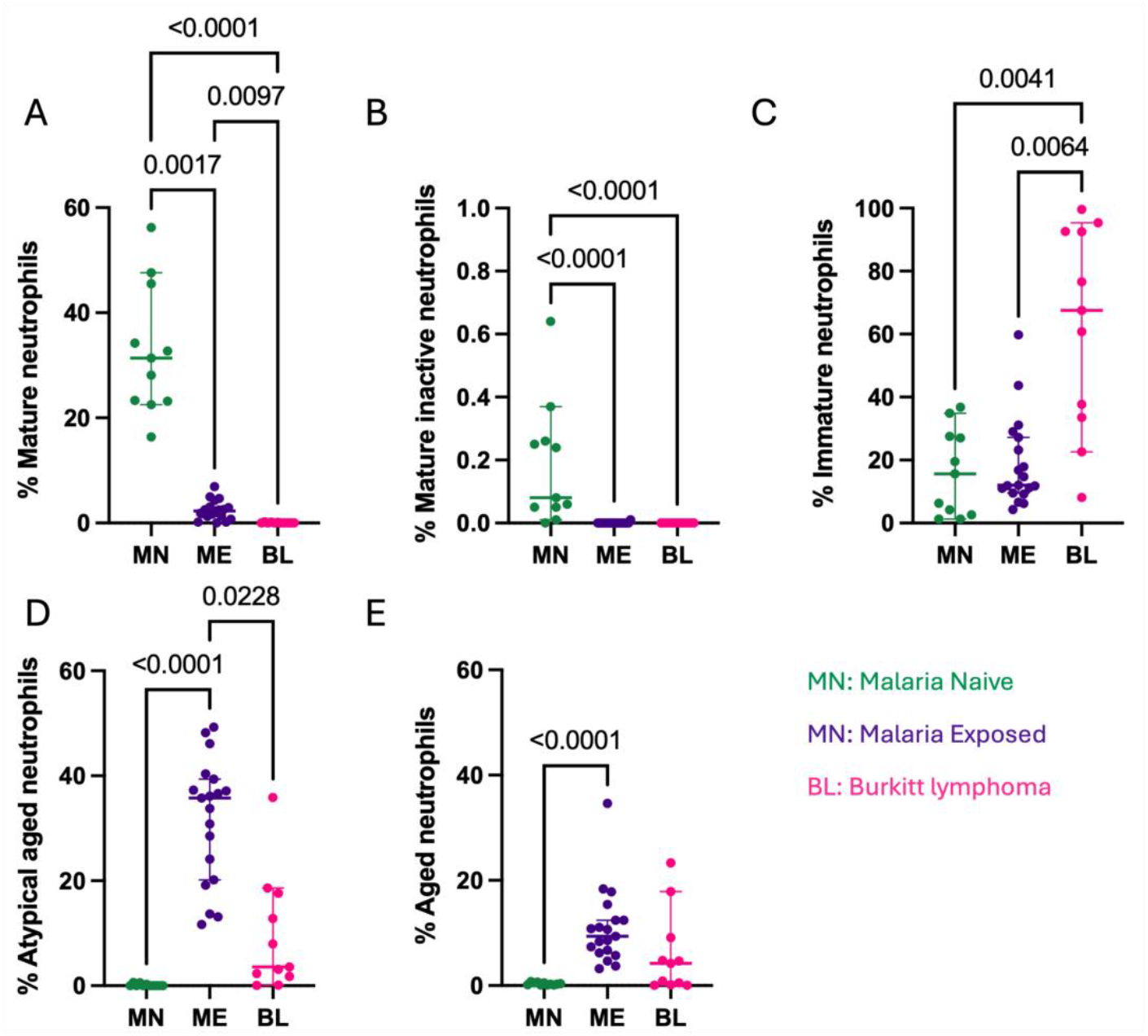
Comparison of neutrophil subset frequency between healthy and BL children. **A**. Mature (CD11b^pos^CD62L^pos^CD182^pos^CD184^neg^), **B**. Mature-inactive (CD11b^pos^CD62L^pos^CD182^pos^ CD184^neg^), **C**. Immature (CD10^neg^**), D**. Aged (CD62L^neg^CD184^pos^), and **E**. Atypical aged (CD62L^pos^CD184^pos^) neutrophils. Malaria-naive (MN, green, n=11), Malaria-exposed (ME, purple, n=19) and children with BL (pink, n=11). The bars represent Median and 95% confidence intervals. Kruskal-Wallis test with Dunn’s correction for multiple comparisons was used, *p*-values <0.05 are indicated in the figure.

### Immature neutrophil is the main subset in the peripheral blood of children with BL

We then compared neutrophil subsets in healthy and BL children. Similarly to what we observed between MN and ME, children with BL showed dramatically lower frequency of the mature subset compared to healthy children with a median of 0.02% of mature neutrophils in BL (range [0%-0.23%]) compared to a median of 2.3% in ME (range [0%-4.97%]); *p=0*.*01*) and a median of 31% in MN (range [16.4%–56.3%]); *p=0*.*002*, **Fig 1A)**. Following the same trends, children with BL had no mature inactive neutrophils leading to a strong statistical difference with their frequencies in MN (*p<0*.*0001*, **Fig 1B**). Interestingly, children with BL had significantly higher frequency of the immature neutrophil subset compared to healthy ME children (median 67.5%, range [8.16%-99.6%] vs median 12%, range [4.29%-59.8%]; *p=0*.*006*, **Fig 1C**). Finally, healthy MEchildren had higher frequency of aged and atypical neutrophils compared to children with BL and this difference was statistically significant for the atypical aged neutrophils (median 35.8%, range [11.7%-49.3%] vs median 3.59%, range [0.07%-35.9%]; *p=0*.*023*, **Fig 1D**). No statistical difference was found in regards to aged neutrophils between ME and BL (**Fig. 1E**). We also created a graphical abstract to show how neutrophil subset frequencies vary in ME, MN, and BL children.

Using Spearman correlation, we determined the relationship between neutrophil subsets in ME and BL children. A significant positive correlation was observed in the immature subset (*rs=0*.*70, p<0*.*0001*). Negative correlations were observed regarding the mature (*rs=-0*.*63, p=0*.*0002)*, atypical aged (*rs=-0*.*70, p<0*.*0001*), aged (*rs=-0*.*32, p=0*.*09*), and mature inactive subsets (*rs=*-0.14, *p<0*.*46*)(**Table 1**).

**Table 1:** Spearman correlation of neutrophil subsets between malaria-exposed children and BL. Neutrophil subsets were defined as: Mature (CD11b^pos^CD62L^pos^CD182^pos^CD184^neg^), Atypical Aged (CD62L^pos^CD184^pos^), Aged(CD62L^neg^CD184^pos^), Immature(CD10^neg^), and Mature Inactive (CD11b^neg^CD62L^pos^CD182^pos^CD184^neg^). Correlation coefficients (rs) and *p*-values were calculated using the Spearman rank test.

| Subsets | Mature | Atypical Aged | Aged | Immature | Mature Inactive |
| --- | --- | --- | --- | --- | --- |
| Spearman's correlation($rs$) | -0.63 | -0.70 | -0.32 | **0.70** | -0.14 |
| *P-value* | $p=0.0002$ | $p<0.0001$ | $p=0.09$ | $p<0.0001$ | $p=0.46$ |
| *95% CI for P* | -0.81 to -0.35 | -0.84 to -0.45 | -0.61 to 0.05 | **0.45 to 0.84** | -0.48 to 0.23 |
| *Sample size* | n=30 | n=30 | n=30 | n=30 | n=30 |

### Cumulative exposure to malaria during childhood drives neutrophil composition to appear similar to adults

There was no difference in the mature subset frequency between adults and ME, however, MN children showed significantly higher frequency than both the adults and ME children (*p*=0.0004, *p*=0.0002, respectively **Fig. 2A**). There was no significant difference in the mature inactive subset frequency between adults and ME, however, MN children had significantly higher frequency than both the adults and ME children (*p*=0.02, *p*<0.0001, respectively **Fig. 2B**). No differences were observed across groups in regards to the immature neutrophil subset (**Fig. 2C**), and there was no significant difference in the aged neutrophil subset frequency between adults and ME (**Fig. 2D**), however, both the adults and ME children had significantly higher frequency of the aged neutrophils compared to the MN children (*p*=0.0003, *p*=0.0002, respectively, **Fig. 2D**). For the atypical aged neutrophil subset, we only observed a significant difference between ME and MN children (*p*<0.0001, **Fig 2E**).

**Figure 2:**
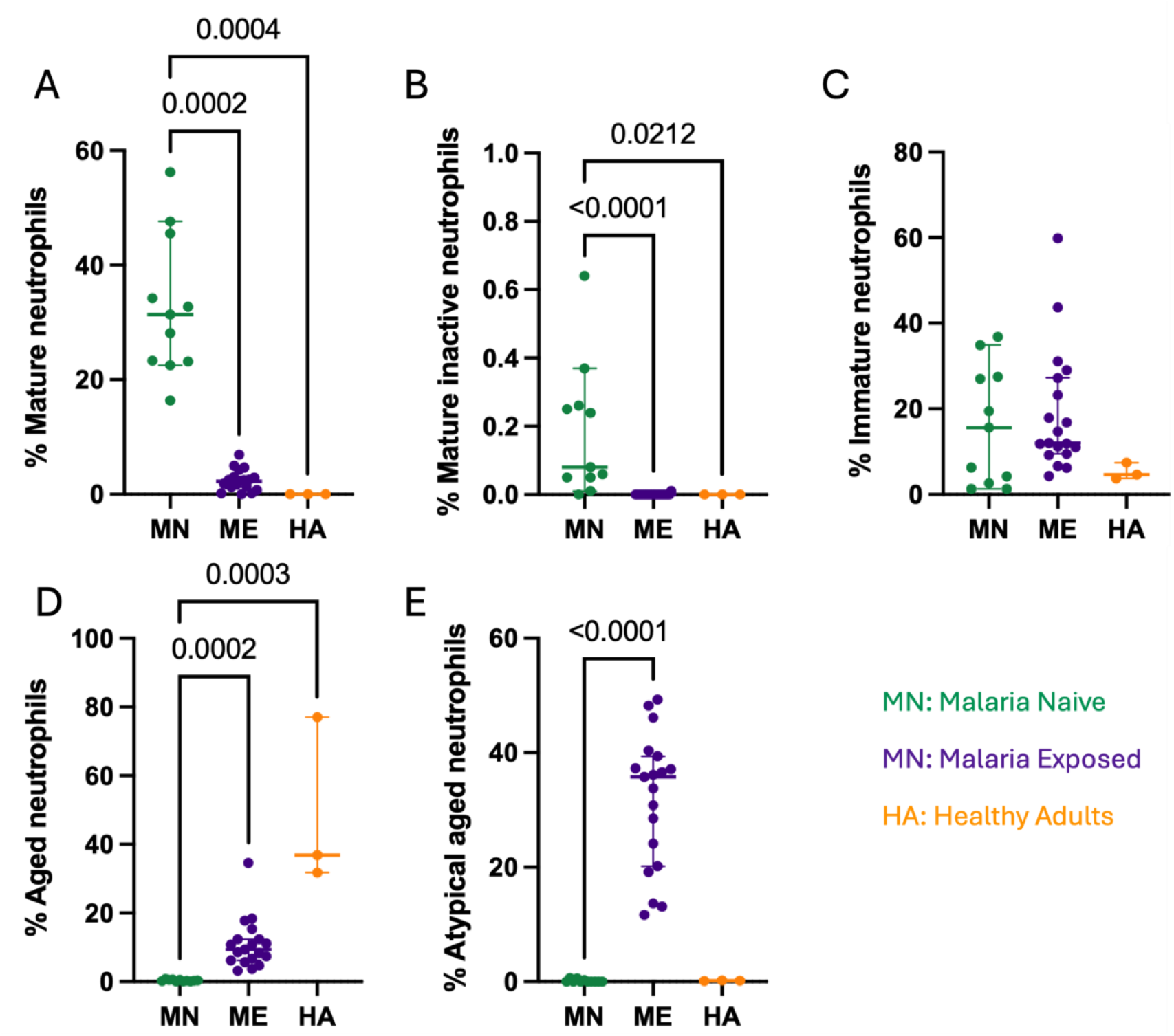
Comparison of neutrophil subset frequency in adults, malaria-exposed children, and malaria-naive children. **A**. Mature (CD11b^pos^CD62L^pos^CD182^pos^CD184^neg^), **B**. Mature-inactive (CD11b^pos^CD62L^pos^CD182^pos^ CD184^neg^), **C**. Immature (CD10^neg^**), D**. Aged (CD62L^neg^CD184^pos^), and **E**. Atypical aged (CD62L^pos^CD184^pos^) neutrophils. Malaria-naive children (MN, green, n=11); Malaria-exposed children (ME, purple, n=19); and healthy adults (HA, orange, n=3). The bars represent Median and 95% confidence intervals. Kruskal-Wallis test with Dunn’s correction for multiple comparisons was used, *p*-values<0.005 are indicated in the plots.

### Children with non-BL cancers have a higher frequency of the atypical aged neutrophil subset

To assess whether the observed neutrophil profiles were unique to BL, we compared them to children with other cancers (OC). No statistical differences were observed in the frequencies of the mature, mature inactive, immature, and aged neutrophil subsets (**Fig. 3A/B/C/D**). The only difference observed was for the atypical aged subset which was higher in the OC compared to BL patients (median of 25.4% atypical aged in OC, range [9.54-65.6] vs median of 3.59% in BL, range [0.07%-35.9%]; *p= 0*.*019*, **Fig. 4E**). These findings suggest that there are immunological changes unique to non-BL cancers.

**Figure 3:**
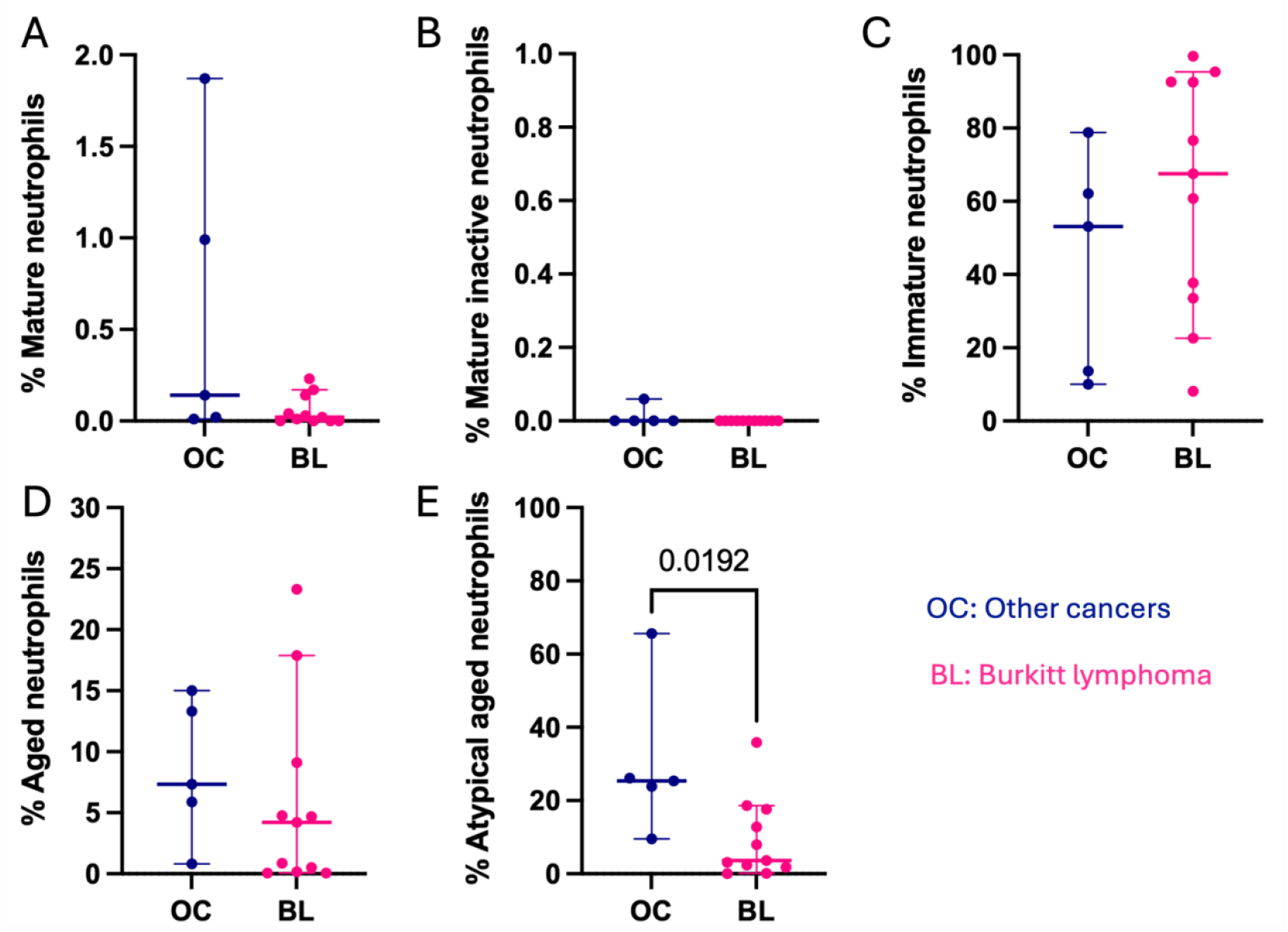
Comparison of neutrophil subset frequency between children with eBL and children with non-BL cancers. **A**. Mature (CD11b^pos^CD62L^pos^CD182^pos^CD184^neg^), **B**. Mature-inactive (CD11b^pos^CD62L^pos^CD182^pos^ CD184^neg^), **C**. Immature (CD10^neg^**), D**. Aged (CD62L^neg^CD184^pos^), and **E**. Atypical aged (CD62L^pos^CD184^pos^) neutrophils. Children with other childhood cancers (OC, blue, n=5); Children with BL (pink, n=11). The bars represent Median and 95% confidence intervals. Mann-Whithney non-parametric test was used. *P*-values<0.05 are indicated in the plots.

## Discussion

Chronic *Pf*-malaria exposure appears to alter proportions of neutrophil subsets by significantly increasing the frequency of the aged and atypical aged subsets, while significantly reducing the frequency of both the mature and mature inactive neutrophil subsets. This shift results in neutrophil profiles that closely resemble those seen in healthy adults, suggesting that chronic malaria exposure drives a shift towards an aged neutrophil phenotype in malaria-exposed children. Additionally, the frequencies of the immature subset in malaria-exposed children and BL children showed a significant correlation.

Chronic malaria exposure has been linked to defects in immunosurveillance, contributing to BL pathogenesis (30,4). Mounting evidence suggests that repeated clinical episodes of *Pf* malaria result in substantial modification of the host immune system, significantly altering the phenotype and function of several immune cell populations (31,32). To understand the effect of repeated malaria exposure on neutrophil subsets, we characterized and compared neutrophil subsets between malaria-exposed and malaria-naive children. The aged neutrophil subset (CD62L^neg^CD184^pos^) and atypical-aged (CD62L^pos^CD184^pos^) frequencies were significantly higher in malaria-exposed children. At homeostasis, aged neutrophils exit circulation and migrate to the bone marrow for elimination (33). However, during acute inflammation, instead of returning to the bone marrow, they rapidly migrate to the inflamed site, leading to elevated levels of aged neutrophils in circulation (19). Given that 62% of our malaria-exposed healthy controls had asymptomatic malaria, which is a chronic, low-grade parasite-driven inflammation, our results suggest that malaria, just like other inflammatory diseases, drives a shift in neutrophil subsets, causing the accumulation of aged neutrophil subsets in circulation.

These malaria-exposed children also had significantly low frequencies of both mature (CD11b^pos^ CD62L^pos^ CD182^pos^ CD184^neg^) and mature-inactive (CD11b^pos^ CD62L^pos^ CD182^pos^ CD184^neg^) subsets. Inflammatory signals and elevated cytokine levels have been shown to modulate the expression of CD182, a critical chemokine receptor that regulates the egress of mature neutrophils from the bone marrow to inflammatory sites (34). This modulation has been linked to a reduced frequency of mature neutrophils in circulation in various inflammatory diseases, such as sepsis (34,35,36), therefore suggesting that the parasite-induced chronic inflammation could be driving the observed low frequency of the mature subsets in malaria-exposed children. In addition to the reduced expression of neutrophil activation marker CD11b previously reported in children with malaria (25), our findings confirm that, similar to other inflammatory diseases, chronic malaria exposure disrupts neutrophil subset homeostasis, leading to the accumulation of aged neutrophil subsets and a concomitant reduction in mature subsets in circulation.

In addition to the aged neutrophil subset (CD62L^neg^CD184^pos^) reviewed in (37), we identified a distinct aged subset expressing CD62L in our participants, which, to our knowledge, has not been previously reported. This finding is particularly intriguing given that aged neutrophils are typically characterized by downregulation of CD62L (38). CD62L, or L-selectin, is a cell adhesion molecule that mediates the multistep neutrophil trans endothelial migration process (39). As neutrophils age, there is downregulation of CD62L and upregulation of CD184 to facilitate their homing to the bone marrow in a rhythmic fashion for efferocytic clearance (40). It is, however, not entirely clear as to whether these phenotypic changes occur at the same rate; this temporal mismatch could lead to a “transient” aged atypical subset. Our observation, therefore, raises the possibility of additional heterogeneity and functional differences within the aged neutrophil compartment that could be relevant to infection-driven and cancer-related immune responses. Further studies are needed to characterize the function and origin of this atypical subset.

Cancer studies have shown that tumors trigger emergency granulopoiesis by producing elevated levels of granulocyte colony-stimulating factor (G-CSF) (41,42). This phenomenon was described as a tumor-induced mechanism that can lead to excessive recruitment of immature neutrophil subsets into the peripheral blood, facilitating immunosuppression and promoting metastasis (43). In our study, we found significantly higher frequency of immature (CD10^neg^) neutrophil subsets in children with BL compared to the malaria-exposed children, suggesting that BL, just like other cancers, induces excessive recruitment of immature neutrophils as a mechanism of immune evasion and progression. Moreover, a strong positive correlation we found regarding the immature neutrophil subset between malaria-exposed and BL children suggests a similar expansion pattern of this subset in both groups. Despite the established link between cancer and immature neutrophils, our observations suggest that chronic or repeated malaria exposure may also be contributing to the elevated presence of immature neutrophils, pointing to a possible immunological bridge between malaria and BL pathogenesis.

We observed that neutrophil subset frequencies are comparable between malaria-exposed children and healthy adults and that both had a significantly high frequency of the aged (CD62L^neg^CD184^pos^) and atypical aged (CD62L^pos^CD184^pos^) subsets. Aging has been shown to directly affect neutrophil homeostasis and functions (44,45). Elderly individuals over the age of 65 have shown accumulation of the aged (CD62L^neg^CD184^pos^) neutrophil subsets in their circulation, playing a role in age-related pathologies such as periodontitis (46). This subset has been shown to have altered migration patterns, enhanced activation, and increased production of reactive oxygen species (ROS), which can contribute to inflammation and tissue damage (47). Our findings align with earlier studies showing that aging and inflammatory diseases, such as malaria disrupts neutrophil subset balance, leading to a high frequency of aged neutrophils in circulation(19). Notably, we observed a significantly higher frequency of the atypical aged subset (CD62L^pos^CD184^pos^) in both healthy adults and malaria-exposed children. The elevated levels of both aged (CD62L^neg^CD184^pos^) and atypical aged (CD62L^pos^CD184^pos^) neutrophils in malaria-exposed children suggest that chronic malaria exposure may drive a shift toward an aged neutrophil phenotype. However, functional assays comparing these subsets in adults and malaria-exposed children are needed to determine whether they share similar immunological roles. We also observed that the healthy adults and malaria-exposed children have a significantly low frequency of both the mature (CD11b^pos^CD62L^pos^CD182^pos^CD184^neg^) and mature inactive (CD11b^neg^CD62L^pos^ CD182^pos^CD184^neg^) neutrophil subsets in circulation. Mature neutrophils circulate in the blood in an inactive form (48), and when they encounter pathogens or inflammatory signals, they become activated (49). The mature active neutrophils are part of the body’s first defence against infections and inflammations (50). Reduced expression of the neutrophil activation marker, CD11b, has been observed in children with severe malaria (25,51). This, therefore, suggests that the low frequency of the mature neutrophil subsets observed reflects a malaria-induced impairment of neutrophil activation in malaria-exposed children. In addition, a significant decline in the release of neutrophils from the bone marrow into the bloodstream in older individuals has been observed (46), and this could be contributing to the low frequency of the mature neutrophil subsets we observed in adults in our study. We acknowledge the small sample size of our healthy adult group, limiting the interpretation of our results.

To assess whether the observed neutrophil subsets were unique to BL and whether their dysregulation was specific to BL or a common feature in childhood cancers. Atypical aged neutrophils (CD62^pos^CD184^pos^) were found to be of higher frequencies in non-BL cancers as opposed to BL, suggesting that neutrophil dysregulation may not be uniform in BL and other childhood cancers. In neutrophils, CD62L is a crucial cell adhesion molecule that facilitates their migration to the site of inflammation (52). Previous studies in cancer have shown that tumors exploit the aged neutrophils to drive inflammation and promote metastasis (53). Therefore, the significantly higher frequency of the atypical aged neutrophils (CD62L^pos^CD184^pos^) observed in our study suggests that non-BL cancers could be inducing the accumulation of aged neutrophils that still express CD62L. However, extensive work is required to functionally distinguish between atypical aged neutrophils still expressing CD62L and aged neutrophils that no longer express CD62L.

## Conclusion

In conclusion, our work successfully demonstrates that chronic malaria exposure reshapes neutrophil composition by increasing the frequency of aged neutrophils while reducing mature neutrophils. Furthermore, the association between immature neutrophils and BL provides insights into how these malaria-driven immune alterations may potentially contribute to BL immuno-pathogolgy.

## Data availability statement

The raw data supporting the conclusion will be made available by the authors without any reservations.

## Acknowledgement

The authors want to thank the KEMRI-UMASS and AMPATH Reference Lab teams at Moi Teaching and Referral Hospital for their incredible work in disease diagnosis and patient care. We also want to thank our study participants for consenting to be part of this study and for giving their samples for this work to be accomplished.

## Author’s contributions and conflict of interest

S.A.: Conceptualization, Investigation, Data curation, Methodology, Formal analysis, Visualization, Original draft preparation

E.A.: Formal analysis,Methodology, Review & Editing

C.O.: Investigation, Resources, Project administration, Review & Editing

T.K.M: Formal analysis, Visualization, Review & Editing

R.K.T: Resources, Review & Editing,Project administration

Festus M. Njuguna: Resources, Project administration

Kibet K. Keitany: Resources, Project administration

Daniel Chepsiror: Resources

C.A.: Supervision, Review & Editing

A.M.M.: Funding acquisition, Project administration, Resources, Review & Editing

A.W.K.: Supervision, Review & Editing, Validation, Methodology

C.S.F.: Conceptualization, Supervision, Review & Editing, Visualization, Validation, Methodology

## The authors declare that they have no conflict of interest

## References

1. Magrath I. Denis burkitt and the African lymphoma. Ecancermedicalscience. 2009 Sep 30;3:159.

2. Hämmerl L, Colombet M, Rochford R, Ogwang DM, Parkin DM. The burden of Burkitt lymphoma in Africa. Infect Agent Cancer. 2019 Aug 1;14:17.

3. Redmond LS, Ogwang MD, Kerchan P, Reynolds SJ, Tenge CN, Were PA, et al. Endemic Burkitt lymphoma: a complication of asymptomatic malaria in sub-Saharan Africa based on published literature and primary data from Uganda, Tanzania, and Kenya. Malar J. 2020 Jul 28;19(1):239.

4. Moormann AM, Bailey JA. Malaria - how this parasitic infection aids and abets EBV-associated Burkitt lymphomagenesis. Curr Opin Virol. 2016 Oct;20:78–84.

5. Moormann AM, Chelimo K, Sumba PO, Tisch DJ, Rochford R, Kazura JW. Exposure to holoendemic malaria results in suppression of Epstein-Barr virus-specific T cell immunosurveillance in Kenyan children. J Infect Dis. 2007 Mar 15;195(6):799–808.

6. Njie R, Bell AI, Jia H, Croom-Carter D, Chaganti S, Hislop AD, et al. The effects of acute malaria on Epstein-Barr virus (EBV) load and EBV-specific T cell immunity in Gambian children. J Infect Dis. 2009 Jan 1;199(1):31–8.

7. Chattopadhyay PK, Chelimo K, Embury PB, Mulama DH, Sumba PO, Gostick E, et al. Holoendemic malaria exposure is associated with altered Epstein-Barr virus-specific CD8(+) T-cell differentiation. J Virol. 2013 Feb;87(3):1779–88.

8. Moormann AM, Snider CJ, Chelimo K. The company malaria keeps: how co-infection with Epstein-Barr virus leads to endemic Burkitt lymphoma. Curr Opin Infect Dis. 2011 Oct;24(5):435–41.

9. Paludan SR, Pradeu T, Masters SL, Mogensen TH. Constitutive immune mechanisms: mediators of host defence and immune regulation. Nat Rev Immunol. 2021 Mar;21(3):137–50.

10. Phillipson M, Kubes P. The Healing Power of Neutrophils. Trends Immunol. 2019 Jul;40(7):635–47.

11. Johansson C, Kirsebom FCM. Neutrophils in respiratory viral infections. Mucosal Immunol. 2021 Jul;14(4):815–27.

12. Adem E, Cruz Cervera E, Yizengaw E, Takele Y, Shorter S, Cotton JA, et al. Distinct neutrophil effector functions in response to different isolates of Leishmania aethiopica. Parasit Vectors. 2024 Nov 11;17(1):461.

13. Ma Y, Zhang Y, Zhu L. Role of neutrophils in acute viral infection. Immun Inflamm Dis. 2021 Dec;9(4):1186–96.

14. Silvestre-Roig C, Hidalgo A, Soehnlein O. Neutrophil heterogeneity: implications for homeostasis and pathogenesis. Blood. 2016 May 5;127(18):2173–81.

15. Ng LG, Ostuni R, Hidalgo A. Heterogeneity of neutrophils. Nat Rev Immunol. 2019 Apr;19(4):255–65.

16. Filep JG, Ariel A. Neutrophil heterogeneity and fate in inflamed tissues: implications for the resolution of inflammation. Am J Physiol Cell Physiol. 2020 Sep 1;319(3):C510–32.

17. Hellebrekers P, Vrisekoop N, Koenderman L. Neutrophil phenotypes in health and disease. Eur J Clin Invest. 2018 Nov;48 Suppl 2(Suppl Suppl 2):e12943.

18. Bongers SH, Chen N, van Grinsven E, van Staveren S, Hassani M, Spijkerman R, et al. Kinetics of neutrophil subsets in acute, subacute, and chronic inflammation. Front Immunol. 2021 Jun 24;12:674079.

19. Uhl B, Vadlau Y, Zuchtriegel G, Nekolla K, Sharaf K, Gaertner F, et al. Aged neutrophils contribute to the first line of defense in the acute inflammatory response. Blood. 2016 Nov 10;128(19):2327–37.

20. Mishra S, Arsh AM, Rathore JS. Trained innate immunity and diseases: Bane with the boon. Clinical Immunology Communications. 2022 Dec;2:118–29.

21. Popa GL, Popa MI. Recent Advances in Understanding the Inflammatory Response in Malaria: A Review of the Dual Role of Cytokines. J Immunol Res. 2021 Nov 8;2021:7785180.

22. Lissner MM, Cumnock K, Davis NM, Vilches-Moure JG, Basak P, Navarrete DJ, et al. Metabolic profiling during malaria reveals the role of the aryl hydrocarbon receptor in regulating kidney injury. Elife [Internet]. 2020 Oct 6;9. Available from: https://pmc.ncbi.nlm.nih.gov/articles/PMC7538157/

23. Aitken EH, Alemu A, Rogerson SJ. Neutrophils and Malaria. Front Immunol. 2018 Dec 19;9:3005.

24. Babatunde KA, Adenuga OF. Neutrophils in malaria: A double-edged sword role. Front Immunol. 2022 Jul 28;13:922377.

25. Mandala WL. Expression of CD11a, CD11b, CD11c, and CD18 on Neutrophils from Different Clinical Types of Malaria in Malawian Children. J Blood Med. 2022 Jan 4;13:1–10.

26. Rainey JJ, Omenah D, Sumba PO, Moormann AM, Rochford R, Wilson ML. Spatial clustering of endemic Burkitt’s lymphoma in high-risk regions of Kenya. Int J Cancer. 2007 Jan 1;120(1):121–7.

27. Mustapha AM, Musembi S, Nyamache AK, Machani MG, Kosgei J, Wamuyu L, et al. Secondary malaria vectors in western Kenya include novel species with unexpectedly high densities and parasite infection rates. Parasit Vectors. 2021 May 12;14(1):252.

28. Epidemiology. FELTP - data quality audit and assessment of epidemic preparedness and response for malaria in Nandi county [Internet]. [cited 2023 Apr 4]. Available from: https://feltp.or.ke/rdqa-and-assessment-of-malaria-epidemic-preparedness-and-response-nandi.html

29. Hofmann N, Mwingira F, Shekalaghe S, Robinson LJ, Mueller I, Felger I. Ultra-sensitive detection of Plasmodium falciparum by amplification of multi-copy subtelomeric targets. PLoS Med. 2015 Mar;12(3):e1001788.

30. Reynaldi A, Schlub TE, Chelimo K, Sumba PO, Piriou E, Ogolla S, et al. Impact of Plasmodium falciparum Coinfection on Longitudinal Epstein-Barr Virus Kinetics in Kenyan Children. J Infect Dis. 2016 Mar 15;213(6):985–91.

31. Bediako Y, Adams R, Reid AJ, Valletta JJ, Ndungu FM, Sodenkamp J, et al. Repeated clinical malaria episodes are associated with modification of the immune system in children. BMC Med. 2019 Mar 13;17(1):60.

32. Moormann AM, Nixon CE, Forconi CS. Immune effector mechanisms in malaria: An update focusing on human immunity. Parasite Immunol. 2019 Aug;41(8):e12628.

33. Ganesh K, Joshi MB. Neutrophil sub-types in maintaining immune homeostasis during steady state, infections and sterile inflammation. Inflamm Res. 2023 Jun;72(6):1175–92.

34. Lee SK, Kim SD, Kook M, Lee HY, Ghim J, Choi Y, et al. Phospholipase D2 drives mortality in sepsis by inhibiting neutrophil extracellular trap formation and down-regulating CXCR2. J Exp Med. 2015 Aug 24;212(9):1381–90.

35. Metzemaekers M, Gouwy M, Proost P. Neutrophil chemoattractant receptors in health and disease: double-edged swords. Cell Mol Immunol. 2020 May;17(5):433–50.

36. Ha H, Debnath B, Neamati N. Role of the CXCL8-CXCR1/2 axis in cancer and inflammatory diseases. Theranostics. 2017 Apr 7;7(6):1543–88.

37. Strydom N, Rankin SM. Regulation of circulating neutrophil numbers under homeostasis and in disease. J Innate Immun. 2013 Apr 6;5(4):304–14.

38. Visan I. Aging neutrophils. Nat Immunol. 2015 Nov 20;16(11):1113–1113.

39. Rahman I, Collado Sánchez A, Davies J, Rzeniewicz K, Abukscem S, Joachim J, et al. L-selectin regulates human neutrophil transendothelial migration. J Cell Sci. 2021 Feb 8;134(3):jcs250340.

40. Adrover JM, Del Fresno C, Crainiciuc G, Cuartero MI, Casanova-Acebes M, Weiss LA, et al. A neutrophil timer coordinates immune defense and vascular protection. Immunity. 2019 Feb 19;50(2):390–402.e10.

41. Mouchemore KA, Anderson RL. Immunomodulatory effects of G-CSF in cancer: Therapeutic implications. Semin Immunol. 2021 Apr;54(101512):101512.

42. Meyer MA, Baer JM, Knolhoff BL, Nywening TM, Panni RZ, Su X, et al. Breast and pancreatic cancer interrupt IRF8-dependent dendritic cell development to overcome immune surveillance. Nat Commun. 2018 Mar 28;9(1):1250.

43. Mackey JBG, Coffelt SB, Carlin LM. Neutrophil Maturity in Cancer. Front Immunol. 2019 Aug 14;10:1912.

44. Li Y, Tian X, Luo J, Bao T, Wang S, Wu X. Molecular mechanisms of aging and anti-aging strategies. Cell Commun Signal. 2024 May 24;22(1):285.

45. Van Avondt K, Strecker JK, Tulotta C, Minnerup J, Schulz C, Soehnlein O. Neutrophils in aging and aging-related pathologies. Immunol Rev. 2023 Mar;314(1):357–75.

46. Wang Z, Saxena A, Yan W, Uriarte SM, Siqueira R, Li X. The impact of aging on neutrophil functions and the contribution to periodontitis. Int J Oral Sci. 2025 Jan 16;17(1):10.

47. Ling S, Xu JW. Phenotypes and functions of “aged” neutrophils in cardiovascular diseases. Biomed Pharmacother. 2024 Oct 1;179(117324):117324.

48. Deniset JF, Kubes P. Neutrophil heterogeneity: Bona fide subsets or polarization states? J Leukoc Biol. 2018 May 1;103(5):829–38.

49. Hong CW. Current understanding in neutrophil differentiation and heterogeneity. Immune Netw. 2017 Oct 16;17(5):298–306.

50. Kraus RF, Gruber MA. Neutrophils-from bone marrow to first-line defense of the innate immune system. Front Immunol. 2021 Dec 23;12:767175.

51. Prah DA, Amoah LE, Gibbins MP, Bediako Y, Cunnington AJ, Awandare GA, et al. Comparison of leucocyte profiles between healthy children and those with asymptomatic and symptomatic Plasmodium falciparum infections. Malar J. 2020 Oct 9;19(1):364.

52. Jafari M, Cardenas EI, Ekstedt S, Arebro J, Petro M, Karlsson A, et al. Delayed neutrophil shedding of CD62L in patients with chronic rhinosinusitis with nasal polyps and asthma: Implications for Staphylococcus aureus colonization and corticosteroid treatment. Clin Transl Allergy. 2024 Mar 10;14(3):e12347.

53. Peng Z, Liu C, Victor AR, Cao DY, Veiras LC, Bernstein EA, et al. Tumors exploit CXCR4hiCD62Llo aged neutrophils to facilitate metastatic spread. Oncoimmunology. 2021 Jan 1;10(1):1870811.

