## Supplemental data for "The Impact of Malaria-Induced Neutrophil Subset Shift and a Link to Burkitt Lymphoma"

**SUPPORTING INFORMATION**

**Flow Cytometric Analysis**

***Reagents****:* The flow cytometry cell staining buffer was prepared with 3% heat-inactivated fetal bovine serum (FBS) in phosphate-buffered saline without calcium chloride and magnesium chloride. (Sigma Life Science #Lot: RNBL4461). Brilliant Staining Buffer (BD Biosciences # Cat: 566349) was used to prevent staining artifacts. Cells were fixed with the 1X Fixation and Permeabilization Solution containing 4.2% formaldehyde (BD Biosciences #Cat:51-2090KZ). UltraComp eBeads Plus (Thermo Fisher Scientific # Lot: 2697000) compensation beads were used for compensation controls.

***Compensation Controls’ staining****:* Appropriately labeled microtubes, including the unstained tube, were used. Approximately 30 μL of compensation beads (UltraComp Beads) from Invitrogen (Thermo Fisher Scientific # Lot: 2697000) was added to each tube. The beads were then stained with 1 μL of the respective antibody and incubated for 30 minutes at room temperature, except for the unstained tube. The beads were washed once in the cell staining buffer. Then fixation was done for 20 minutes in 250 μL of BD cytofix/cytoperm, a fixation/permeabilization solution with 4.2% formaldehyde at 4°C, followed by washing in a cell staining buffer. After centrifugation and removing the supernatant, beads were resuspended in 300 μL of flow cytometry staining buffer containing sodium azide. They were then kept at 4°C overnight for analysis the following day, together with the samples.

***Sample and Fluorescence Minus One (FMO) controls staining:*** Whole blood samples were taken from one individual throughout the study for all the runs. Appropriately labeled microtubes were used for the required FMOs and samples. 100 μL of the control sample was aliquoted into each FMO tube. 100 μL of each sample was also aliquoted into an appropriately labelled sample tube. Live-dead staining was performed on all samples, plus the FMOs. Live-dead staining was done by adding 100 μL of 1 /1000 dilution of viability dye, followed by a 15-minute incubation in the dark. The prepared FMO cocktails and sample antibody cocktails were added to FMO and sample tubes, respectively. The tubes were incubated for 30 minutes at 4 °C in the dark. After incubation, the cells were fixed in 250 μL of fixation/permeabilization solution with 4.2% formaldehyde for 20 minutes at 4°C and washed twice with 1 mL cell staining buffer. The cells were then resuspended in 500 μL of Flow Cytometry Staining Buffer (containing Sodium Azide) and kept at 4°C overnight. The following day, the samples and FMOs were filtered into appropriately labeled flow tubes using a 70 μm cell filter at room temperature. The compensation controls were also transferred to the flow tubes before analysis in the flow cytometer. Data was acquired in the Beckman Coulter flow cytometer using the CytoFLEX software.

**Supplementary Figures**

**S1 FIGURE: S1 FigA:** The flow cytometry gating strategy used for phenotyping neutrophils and Boolean gating used to identify neutrophil subsets

**S1 FIGURE: S1 Fig B**: Gating strategy for CD62L FMO, malaria-exposed children, malaria-naive children, Adults, and children with other childhood cancers

**S1 FIGURE: S1 Fig C**: **Gating strategy for identifying atypical neutrophil subset**.

FMO used to gate for CD184 and CD62L is also included. The arrows are pointing in the direction of the workflow.

**Supplementary Tables**

**S1 TABLE: Neutrophil flow cytometry antibody panel.** The table lists the markers used to identify neutrophils, the antibodies against the markers, the fluorochrome on the antibodies, the antibody clone, and the antibody RRIDs

| **Antibody** | **Fluorochrome** | **Clone** | **RRID** |
| --- | --- | --- | --- |
| CD3 | Alexa Fluor700 | OKT3 | AB_2563408 |
| CD19 | Alexa Fluor700 | HIB19 | AB_2616936 |
| CD56 | Alexa Fluor700 | 5.1H11 | AB_2564099 |
| CD15 | PerCP | HI98 | AB_893256 |
| CD16 | BV510 | 3G8 | AB_2562085 |
| CD10 | FITC | HI10a | AB_314919 |
| CD11b | BV605 | ICRF44 | AB_256202 |
| CD62L | BV421 | DREG-56 | AB_2562914 |
| CD182 | APC | 5E8-C7-F10 | AB_492936 |
| CD184 | PE | 12G5 | AB_314612 |

**S2 TABLE: Demographics and Characteristics of the Study Participants. Mal-exposed HC**: Malaria-exposed healthy controls. **Mal naive HC**: Malaria-naive healthy controls. **BL**: endemic Burkitt's Lymphoma cases. **WBC**: White Blood Cell Count. **ANC**: Absolute Neutrophil Count. **^A^**: Median [min-max]. **^P^**: p-value from the Mann-Whitney statistical test. **^C^**: p-value from the Chi-square statistical test. **Hgb**: Hemoglobin. **qPCR**: Malaria parasite DNA quantified by quantitative Polymerase Chain Reaction. **AMA1**: antibodies against Apical Membrane Antigen 1. Significant p-values are in bold.

|  | Mal exposed HC (n=19) | Mal naive HC (n=11) | BL Cases   (n=11) | Other Cancer Cases  (n = 5) | Mal exposed vs. Mal naive | Mal exposed vs. BL | Mal naive vs. BL | BL vs. Other Cancers |
| --- | --- | --- | --- | --- | --- | --- | --- | --- |
| Age^A^  (years) | 5  [4-7] | 5  [4-7] | 6  [3-9] | 8  [6-14] | 0.71^P^ | **0.03**^P^ | 0.23^P^ | 0.07^P^ |
| Sex  (% of males) | 52.6% | 36.3% | 81% | 60% | 0.4^C^ | 0.11^C^ | **0.33**^C^ | 0.37^C^ |
| ANC^A^  (10^3^/µL) | 32.35  [18-69] | 40.2  [26.1-60.5] | 58.4  [37.6-76.2] | 74.75  [60.5-87.5] | 0.09^P^ | **0.0001**^P^ | **0.02**^P^ | **0.04**^P^ |
| WBC^A^  (10^3^/µL) | 6.1  [3.1-9.2] | 7.22  [5.48-8.28] | 8.26  [5.1-13.28] | 6.49  [4.12-8.15] | 0.05^P^ | **0.01**^P^ | 0.18^P^ | 0.18^P^ |
| qPCR  (% positive) | 68% |  | 1% | 1% |  |  |  |  |
| Hgb  (g/dL)^A^ | 12.3  [10.6-14.1] | 13.7  [11.8-15.7] | 8.7  [7-17] | 9.2  [8-14] | **0.01**^P^ | **0.002**^P^ | **0.01**^P^ | 0.69^P^ |
| AMA1^A^  (MFI) | 72126  [384-217702] | 0  [0-2541] | 2495  [0-171811] | 197  [0-153896] | **0.0001**^P^ | 0.10^P^ | **0.01**^P^ | 0.27^P^ |

**S3 TABLE**: **Demographics and Characteristics of the Adult Participants. WBC**: White Blood Cell Count. **ANC**: Absolute Neutrophil Count. **^A^**: Median [min-max].**Hgb**: Hemoglobin. **qPCR**: Quantitative Polymerase Chain Reaction. **AMA1**: Apical Membrane Antigen 1.

|  | Age^A^  (years) | Sex  (% of males) | ANC^A^  (10^3^/µL) | WBC^A^  (10^3^/µL) | Hgb^A^  (g/dL) | AMA1^A^  (MFI) |
| --- | --- | --- | --- | --- | --- | --- |
| Healthy Adults (n=3) | 29  [34-26] | 100% | 52.9  [66.2-35.9] | 5.24  [5.712-5.13] | 16  [16-15.5] | 31,108  [59775-104] |

**S4 TABLE: Frequency of neutrophil subsets in the study populations.** Frequency **^A^**: Median [min-max].

| Study population | **Mature**  **(CD11b+ CD62L+ CD182+ CD184-)**  ***[****Frequency^A^****]*** | **Atypical Aged**  **(CD62L+ CD184+)**  ***[****Frequency^A^****]*** | **Aged**  **(CD62L- CD184+**)  ***[****Frequency^A^****]*** | **Mature Inactive (CD11b- CD62L+ CD182+ CD184-)**  ***[****Frequency^A^****]*** | **Immature (CD10)**  ***[****Frequency^A^****]*** |
| --- | --- | --- | --- | --- | --- |
| Malaria-  Exposed | 2.3^A^  [0-4.97] | 35.8^A^  [11.7-49.3] | 9.36^A^  [3.2-34.6] | 0 | 12^A^  [4.29-59.8] |
| Malaria-  naive | 31.4^A^  [16.4-56-3] | 0.02^A^  [0.02-0.64] | 0.3^A^  [0.15-0.78] | 0.08^A^  [0-0.37] | 15.6^A^  [1.28-36-8] |
| BL | 0.02^A^  [0-0.23] | 3.59^A^  [0.07-35.9] | 4.22^A^  [0.05-23.3] | 0 | 67.5^A^  [8.16-99.6] |
| Other cancers | 0.14^A^  [0.02-1.87] | 25.4^A^  [9.54-65.6] | 7.33^A^  [0.82-15] | 0 | 53.1^A^  [10-78.8] |
| Adults | 0.01^A^  [0-0.02] | 0.19^A^  [0.15-0.22] | 36.9^A^  [31.8-77] | 0 | 4.6^A^  [3.75-7.46] |
